# Automatic vocalisation detection delivers reliable, multi-faceted, and global avian biodiversity monitoring

**DOI:** 10.1101/2023.09.14.557670

**Authors:** Sarab S. Sethi, Avery Bick, Ming-Yuan Chen, Renato Crouzeilles, Ben V. Hillier, Jenna Lawson, Chia-Yun Lee, Shih-Hao Liu, Celso Henrique de Freitas Parruco, Carolyn Rosten, Marius Somveille, Mao-Ning Tuanmu, Cristina Banks-Leite

## Abstract

Tracking biodiversity and its dynamics at scale is essential if we are to solve global environmental challenges. Detecting animal vocalisations in passively recorded audio data offers a highly automatable, inexpensive, and taxonomically broad way to monitor biodiversity. However, uptake is slow due to the expertise and labour required to label new data and fine-tune algorithms for each deployment. In this study, we applied an off-the-shelf bird vocalisation detection model, BirdNET, to 152,376 hours of audio comprising of datasets from Norway, Taiwan, Costa Rica, and Brazil. We manually listened to a subset of detections for each species in each dataset and found precisions of over 80% for 89 of the 139 species (100% for 57 species). Whilst some species were reliably detected across multiple datasets, the performance of others was dataset specific. By filtering out unreliable detections, we could extract species and community level insight on diel (Brazil) and seasonal (Taiwan) temporal scales, as well as landscape (Costa Rica) and national (Norway) spatial scales. Our findings demonstrate that, with a relatively fast validation step, a single vocalisation detection model can deliver multi-faceted community and species level insight across highly diverse datasets; unlocking the scale at which acoustic monitoring can deliver immediate applied impact.

## Introduction

Biodiversity plays a crucial role in food security, disease dynamics, human wellbeing, and more^1^. However, the vast number of species and the complexity of their interactions makes tracking biodiversity and understanding how it is impacted by anthropogenic activities a challenge^2^. Reliable, scalable, and taxonomically diverse biodiversity monitoring is therefore essential if we are to thrive sustainably as a society.

Traditional biodiversity surveys are time consuming and rely on niche expertise. However, recent declines in cost and increased accessibility of robotics platforms and electronic sensors have transformed our ability to survey ecosystems at larger scales^3^. Using autonomous sensing technologies, scientists have tracked cetaceans in the Pacific from drones^4^, mammals in the Serengeti with camera traps^5^, and bats across London from ultrasonic microphones^6^. However, the machine learning models used in each of these cases were trained on manually labelled subsets of data from the study system of interest. “Plug-and-play” approaches which convert raw field sensor data into reliable species community insight across diverse ecosystems, without retraining models, have not been demonstrated to work reliably to date.

Due to the diversity of species and their behaviours, we are unlikely to ever develop a single technology to monitor all biodiversity in all ecosystems (there are no free lunches^7^). Nevertheless, detection and classification of animal vocalisations in long-term acoustic recordings is a promising approach in its ability to scale well temporally, spatially, and taxonomically. Bird vocalisations in particular have been recorded by hobbyists and scientists for decades culminating in rich libraries of annotated data which span the globe^8^. Classification models have been trained on these libraries^9,10^ and some studies have evaluated the performance of these models on single datasets^11^. However, no studies have looked at the performance of vocalisation classification models when applied to multiple large acoustic datasets collected across diverse ecosystems.

In this study we investigated the following open questions: (i) How reliably can we monitor bird communities in diverse ecosystems across the globe using a single state-of-the-art vocalisation classification model, and (ii) what types of biodiversity insight could such an approach deliver?

## Results

We collected 152,376 hours of passively recorded acoustic data from temperate forests across Norway (76,746 hours), tropical and subtropical forests across Taiwan (49,548 hours), diverse tropical landscapes across the Osa Peninsula in Costa Rica (25,305 hours), and tropical forests in the Amazon, State of Para, Brazil (777 hours) (**Fig. 1a, 1e**). We used BirdNET^9^, the largest open-source vocalisation classification neural network model available, to automatically extract 627,995 bird vocalisations from 379 species from the audio data (**Fig. 1b**). To ensure that we had enough data for verification, we only considered species with over 50 detections in each dataset, giving a filtered list of 625,063 detections from 135 species (**Fig. 1c**).

**Figure 1:**
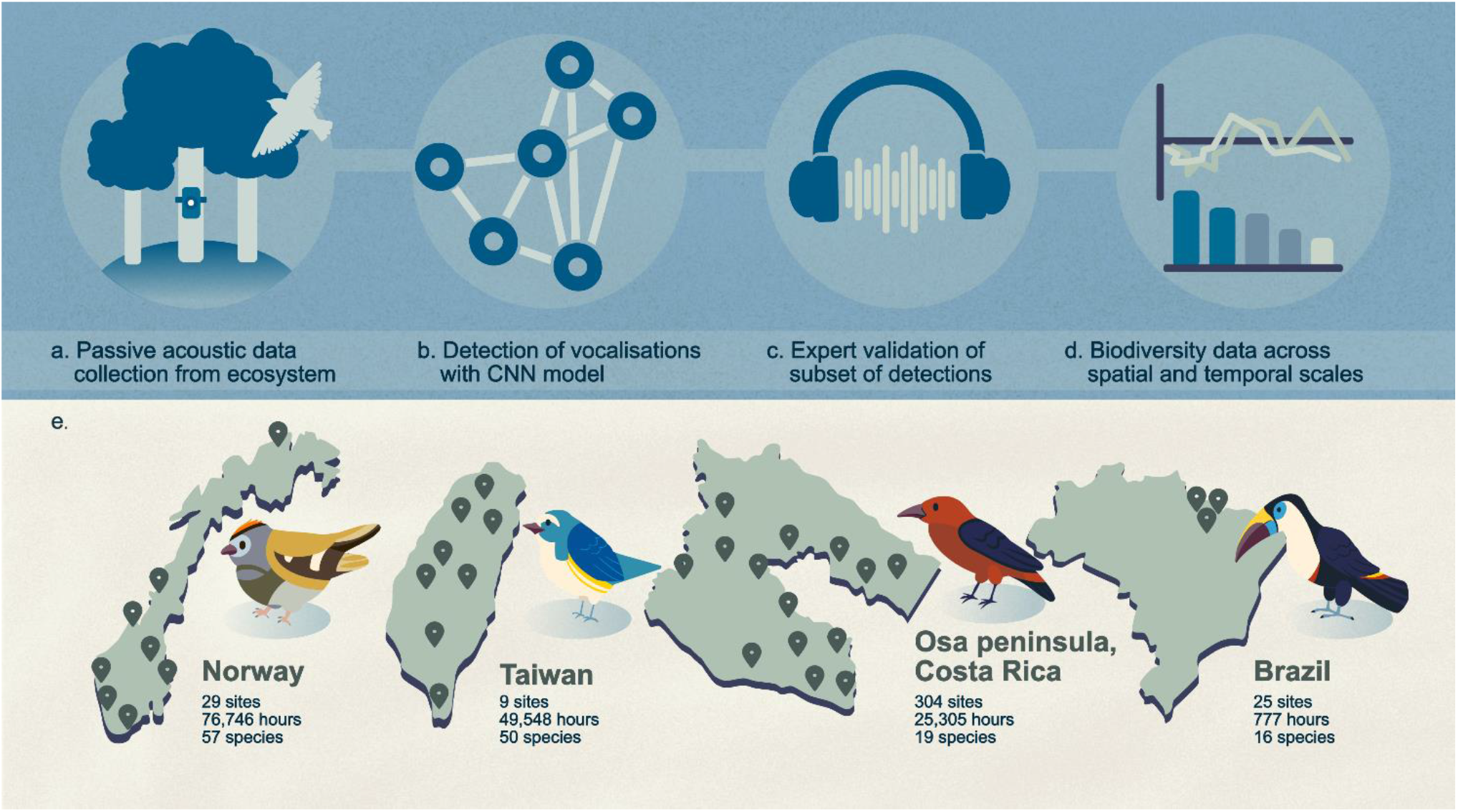
Schematic overview of the study. **(a)** We deployed passive acoustic monitoring devices in ecosystems to collect large scale data. **(b)** The audio data was run through BirdNET, a state-of-the-art convolutional neural network model, to automatically detect and classify bird vocalisations. **(c)** We manually labelled a subset of the detections for each species in each dataset. **(d)** We used detections (filtered to only include well-performing species) to derive reliable species and community insight across a variety of spatial and temporal scales. **(e)** Map showing approximate locations of surveys sites across Norway, Taiwan, the Osa Peninsula in Costa Rica, and State of Para in Brazil. Species depicted are Goldcrest (Norway), Red-flanked Bluetail (Taiwan), Scarlet-rumped Tanager (Costa Rica), and White-throated Toucan (Brazil).

We listened to 50 randomly chosen detections for each species from each dataset, and labelled them as correct, unsure, or incorrect to quantify the precision of the model (c. 20-30 minutes labelling effort per species, **Fig. 1c**). Measuring recall for so many species was intractable on such large datasets^11^. In Norway 25/57, Taiwan 14/40, Costa Rica 11/19, and Brazil 7/16 species all had 100% precision – i.e., every vocalisation listened to for the given species was correct (**Fig. 2**). Allowing for precision rates of 80% or above further increased the number of species that could be considered accurately detectable with the model (41/57 Norway, 18/40 Taiwan, 19/19 Costa Rica, 14/16 Brazil).

**Figure 2:**
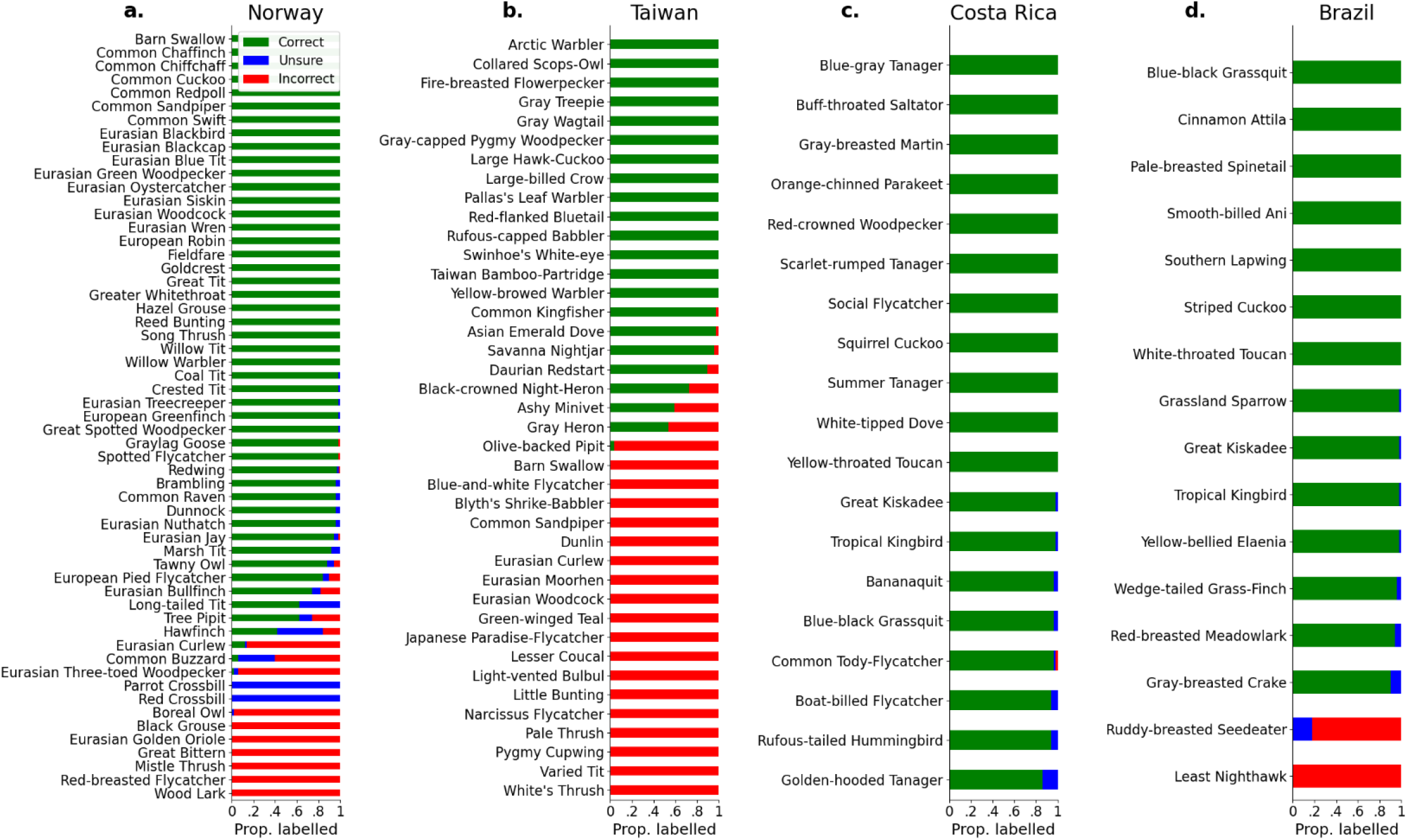
BirdNET was highly precise for many species across diverse datasets. We used BirdNET to automatically extract vocalisations from the audio. 50 detections of each species in each dataset were listened to manually to measure the model’s precision. **(a)** 41/57 species in Norway, **(b)** 18/40 in Taiwan, **(c)** 19/19 in Costa Rica, and **(d)** 14/16 in Brazil all had over 80% precision.

Seven species appeared in two datasets. Blue-black Grassquits, Great Kiskadees, and Tropical Kingbirds had precisions of over 80% in both Brazil and Costa Rica datasets. Detections of Barn Swallows, Eurasian Woodcocks, and Common Sandpipers were reliable in Norway, but not Taiwan. Detections of the Eurasian Curlew were unreliable in both Norway and Taiwan. The inconsistent precision of the same species across datasets might be explained by varying dialects, microphones, experts performing the labelling, geophony, anthropophony, and more, indicating that model performance must be re-characterised for each new deployment of acoustic sensors.

Considering only detections of species with 100% precision, we were able to extract biodiversity insight on small and large spatial and temporal scales, and at both species and community levels (**Fig. 3**). In the Brazilian Amazon, diel vocal activity varied between species which were more vocal at dawn (e.g., Pale-breasted Spinetail), during the day (e.g., Blue-black Grassquit), and at dusk (e.g., White-throated Toucan) (**Fig. 3a**). On the Osa Peninsula in Costa Rica, we found species diversity to be higher in grasslands, a result likely driven by the selection of very common generalist species that were detected reliably by the vocalisation detection model (**Fig. 3b**). Across the temperate forests of Norway, we saw the northward movement of the migratory Willow Warbler throughout spring (**Fig. 3c**). In the forests of Taiwan, we found complex species occurrence patterns across a two-year period, from species which visited Taiwan to breed (e.g., Large Hawk-Cuckoo), those which wintered in the country (e.g., Yellow-browed Warbler), to those which were endemic and vocalising year-round (e.g., Taiwan Bamboo-Partridge) (**Fig. 3d**). Whilst we have showcased known phenomena, we also provide the full list of reliable detections in each dataset for others to explore and investigate new scientific questions in the accompanying supplementary material.

**Figure 3:**
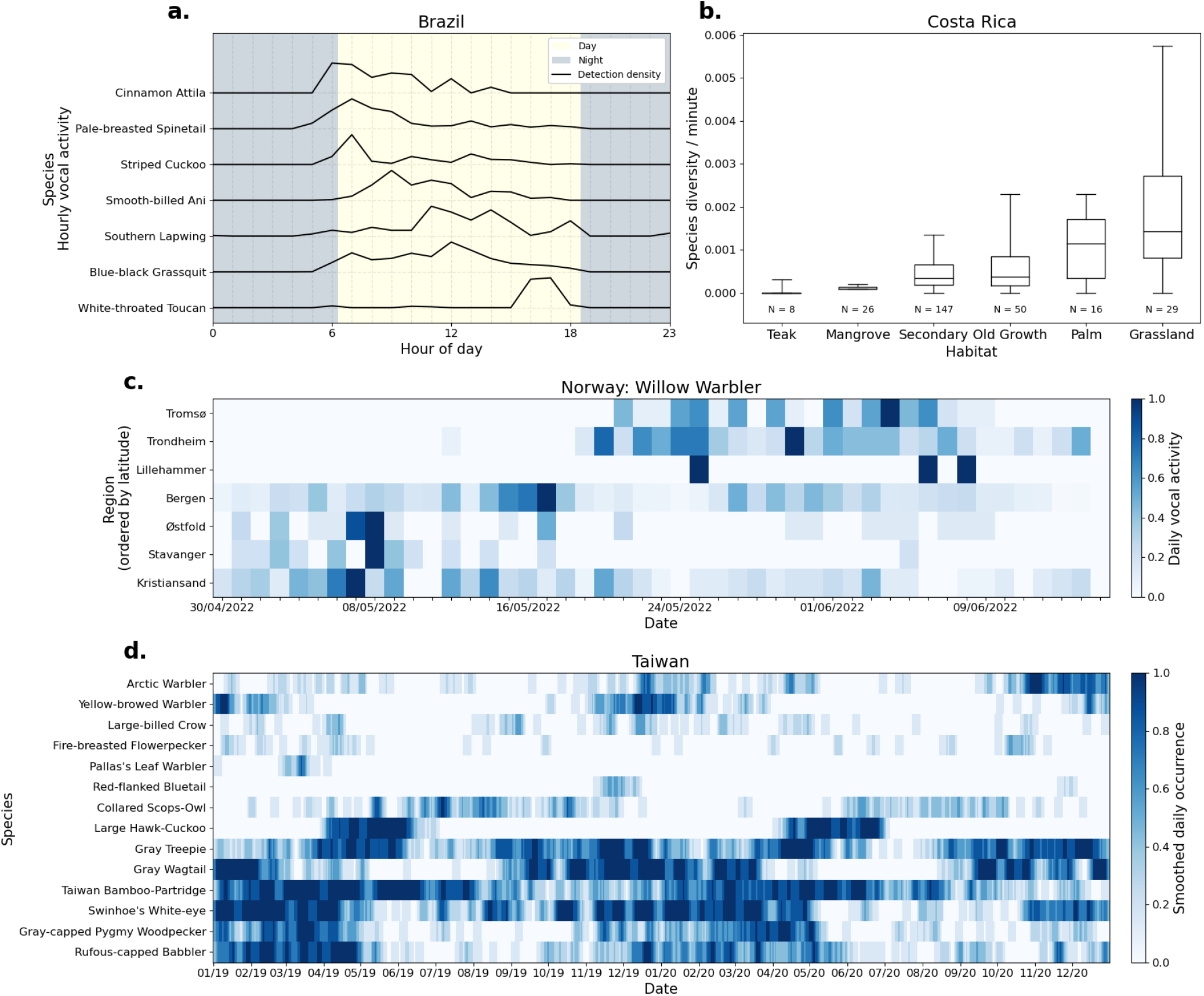
Vocalisation detection data delivered both species and community insight at a range of spatial and temporal scales. **(a)** In Brazil diel detection patterns varied between species which were more vocal at dawn, throughout the day, and at dusk. **(b)** On the Osa Peninsula in Costa Rica, species diversity per minute was heavily influenced by habitat type. Whiskers show 5^th^ and 95^th^ percentiles, and boxes show 25^th^, 50^th^ (i.e., median), and 75^th^ percentiles of the species diversity data. **(c)** In Norway, the Willow Warbler’s northward passage through the country in spring was reconstructed from daily vocal activity in each region. **(d)** Across two years of monitoring in Taiwan, we saw complex dynamics from migratory species which breed or winter in the country, as well as those which were endemic and vocalised throughout the year. Species are ordered using a hierarchical clustering approach.

## Conclusion

We demonstrated that a single vocalisation classification model could deliver reliable monitoring for many bird species across four large datasets spanning diverse habitats. Whilst predictions were not perfect and performance was biased towards common species, with the relatively fast validation of a subset of detections, fine resolution and taxonomically broad biodiversity insight could be unlocked on a variety of temporal and spatial scales. As training datasets grow in size and accessibility and include other animal groups (e.g., fish^12^, insects^13^), the potential applications for vocalisation detection models will widen further. Combined with autonomous data collection approaches^14^, our results pave the way for fully automated acoustic monitoring to be deployed at scale and deliver immediate impact around the world^15^.

## Materials and Methods

### Acoustic data

Norway: We surveyed 29 sites in seven regions across the country between March and October 2022. Sites were all in temperate forests, and devices were spaced at least 100m apart. 76,746 hours of single channel audio was recorded continuously at 44.1kHz using Bugg devices^14^ (with Knowles SPH0641LU4H-1 MEMS microphones) mounted between 1-2m from ground level.

Taiwan: We surveyed 9 sites across the country between January 2019 and December 2020. Sites were all in broadleaf forests and spaced at least 17km apart. 49,548 hours of stereo audio was recorded on a schedule of 1 minute every 3 minutes for 24 hours at 44.1kHz using Wildlife Acoustics Song Meter 4 devices mounted between 1-2m from ground level.

Costa Rica: We surveyed 304 sites on the Osa Peninsula between December 2018 and August 2019. Sites were in varying habitats and spaced at least 500m apart. 25,305 hours of single channel audio was recorded continuously between 05:30 and 9:30, 14:30 and 18:30 and 21:00 and 03:00 at 48,000 kHz using Audiomoth^16^ devices mounted between 1-2m from ground level.

Brazil: We surveyed 25 sites municipality of Mãe do Rio, in the Para State in February 2022. Sites were in located in riparian forests and pastures and spaced at least 500m apart. 777 hours of single channel audio was recorded on a schedule of 1 minute every 5 minutes for 24 hours at a sample rate of 48 kHz using 25 devices mounted between 1-2m from ground level.

### BirdNET model

We used a convolutional neural network (CNN) model, BirdNET, to detect bird vocalisations^9^. For Norway, we used BirdNET-Lite (https://github.com/kahst/BirdNET-Lite) accessed in November 2022. For Taiwan, Costa Rica, and Brazil, we used the newer BirdNET-Analyzer model v2.3 (https://github.com/kahst/BirdNET-Analyzer) through the birdnetlib Python library. Re-analysing the Norway data with BirdNET-Analyzer would have been ideal, but we lacked the resources to relabel the randomly selected detections, and the conclusions of our study were unlikely to have changed. Location data was provided to BirdNET to filter for only species expected at each recorder (based on eBird observations).

To acquire valid BirdNET inputs we resampled audio to 48kHz and generated non-overlapping three-second spectrograms (window size 8ms, 25% overlap, 64 Mel-scaled frequency bands). BirdNET outputs with a confidence of over 0.8 were denoted as detections. In Norway, adjacent detections of the same species were joined together, but in other datasets each three-second detection was kept independently. We only considered species with at least 50 detections within each dataset.

### Measuring precision

For each species in each dataset, we randomly selected 50 detections. Experts familiar with each study region listened to the randomly selected detections and labelled each BirdNET prediction as correct, incorrect, or unsure (e.g., if the call was ambiguous). To measure precision, we only counted the correct labels as true positives.

### Species and community analyses

For the following analyses, we only considered detections from species with 100% precision in Fig. 2.

Brazil: We binned detections by hour of day to measure diel patterns in vocal activity. To normalise the data shown in Fig. 3a, for each species, we divided hourly numbers of detections by the maximum number of detections in a one-hour period.

Costa Rica: For each day at each site, we counted the number of unique species detected and normalised for sampling effort by dividing by the number of minutes recorded on that day. This produced a rudimentary measure of species diversity per minute. Then, to produce Fig. 3b, we separated sites by habitat type using land use maps provided by NASA, created at a scale of 5 × 5 m using Landsat 5 Thematic Mapper (TM) and Landsat 8 Operational Land Imager (OLI).

Norway: We focused on detections of the Willow Warbler between 30/04/2022 and 15/06/2022, and sites were aggregated by region. To show temporal variations in relative vocal activity across regions in Fig. 3c, detections were binned by day, and we divided daily numbers of detections by the maximum number of detections in a day for each region.

Taiwan: We combined detections across all sites, and derived daily occurrence (presence/absence) data for each species. For Fig. 3d, we smoothed the daily occurrence data by convolving the binary occurrence time series for each species with a vector of ones of shape [1x7] (i.e., at the weekly resolution).

## Data Availability

Code to reproduce figures and results presented in this manuscript can be found at https://doi.org/10.5281/zenodo.8338721, with data required to run the code hosted at https://doi.org/10.5281/zenodo.8340251.

## Acknowledgements

We would like to thank the following for their assistance in collecting and processing data; Norway: Tom Roger Østerås, NINA field technicians; Taiwan: the Medium Altitude Experimental Station of Taiwan Endemic Species Research Institute, Dongyanshan Forest Recreation Area, Fonghuanggu Bird and Ecology Park, Liang-Fang Zhang at Wucheng Elementary School.

The project was supported by the following funding sources; Taiwan: National Science and Technology Council (NSTC 111-2321-B-002-019 and 111-2927-I-001-513), Biodiversity Research Center at Academia Sinica, A Royal Society International Exchanges 2021 Cost Share (MOST) grant (Ref: IEC\R3\213038); Norway: Miljødirektoratet; Costa Rica: Natural Environment Research Council (grant no. NE/L002515/1).

Permits were issued by the government of Costa Rica for this study as below: no. de resolución (permiso de investigación)/research permit no.: ACOSA-INV-067-18.

